# How fast are viruses spreading in the wild?

**DOI:** 10.1101/2024.04.10.588821

**Authors:** Simon Dellicour, Paul Bastide, Pauline Rocu, Denis Fargette, Olivier J. Hardy, Marc A. Suchard, Stéphane Guindon, Philippe Lemey

## Abstract

Genomic data collected from viral outbreaks can be exploited to reconstruct the dispersal history of viral lineages in a two-dimensional space using continuous phylogeographic inference. These spatially explicit reconstructions can subsequently be used to estimate dispersal metrics allowing to unveil the dispersal dynamics and evaluate the capacity to spread among hosts. Heterogeneous sampling intensity of genomic sequences can however impact the accuracy of dispersal insights gained through phylogeographic inference. In our study, we implement a simulation framework to evaluate the robustness of three dispersal metrics — a lineage dispersal velocity, a diffusion coefficient, and an isolation-by-distance signal metric — to the sampling effort. Our results reveal that both the diffusion coefficient and isolation-by-distance signal metrics appear to be robust to the number of samples considered for the phylogeographic reconstruction. We then use these two dispersal metrics to compare the dispersal pattern and capacity of various viruses spreading in animal populations. Our comparative analysis reveals a broad range of isolation-by-distance patterns and diffusion coefficients mostly reflecting the dispersal capacity of the main infected host species but also, in some cases, the likely signature of rapid and/or long-distance dispersal events driven by human-mediated movements through animal trade. Overall, our study provides key recommendations for the lineage dispersal metrics to consider in future studies and illustrates their application to compare the spread of viruses in various settings.

## Introduction

Unravelling the spatio-temporal dynamics of pathogenic spread constitutes a long-standing challenge in epidemiology. When designing and implementing intervention strategies to mitigate outbreaks or endemic circulation of pathogens, evaluating the speed at which they spread and circulate within host populations can be crucial. While geo-localised infectious cases can be used to model and quantify the wavefront progression of an outbreak during its expansion phase (see e.g. [1,2]), their analysis usually provides little information about the routes taken by the underlying transmission chains. Genomic sequencing of fast-evolving pathogens however offers the possibility to infer evolutionary relationships among sampled cases. Estimated through a phylogenetic tree, such inferred evolutionary links can for instance provide key insights into the dispersal history and dynamics of a transmission chain [3]. This can be achieved through phylogeographic inference, either performed in discrete [4–6] or continuous space [7,8], which basically consists in mapping time-scaled phylogenetic trees of fast-evolving organisms such as RNA viruses (see e.g. [9,10]) or, sometimes, DNA viruses (see e.g. [11,12]) or bacteria (see e.g. [13,14]).

In particular, continuous phylogeographic inference allows estimating a spatially explicit reconstruction of the dispersal history of the lineages leading to the sampled genomic sequences. The reconstructions in two-dimensional space can subsequently be exploited to gain valuable information about the dispersal dynamics of circulating lineages [15], including their capacity to disperse in space and time. However, for a given outbreak, phylogeographic reconstructions are generally conducted on genomic sequences from a limited proportion of cases, with sampling intensities that can vary considerably among studies.

In the context of these heterogeneous sampling efforts, the goal of the present study is to assess the usefulness of dispersal metrics to evaluate and compare the dispersal dynamic of viral lineages. Specifically, we first aim to assess the robustness of three metrics — a lineage dispersal velocity, a diffusion coefficient, and an isolation-by-distance (IBD) signal metric — to the sampling intensity, i.e. the number of genomic samples considered in the phylogeographic analysis. In addition, we aim to exploit the metrics identified as robust to the sampling effort to compare the dispersal capacity across a variety of viruses of public and one health importance (e.g., Lassa, rabies, avian influenza, or West Nile viruses).

At relatively large scales, dispersal dynamics of viruses primarily spreading within human populations are generally better captured by discrete diffusion processes involving the rapid exchange of viral lineages among remote locations — a typical pattern for remote locations mainly connected within the air traffic network (see e.g. [16,17]). This is for instance the case for the human influenza virus H3N2, whose dispersal at the global scale has been demonstrated to be better predicted by international air travel than geographic distances [18]. In our study, we focus on viruses primarily circulating in animal populations to target dispersal processes that are relatively strongly dependent on geographic distance, and which can by essence be modelled through a continuous diffusion process, such as the relaxed random walk (RRW) diffusion model implemented in the software package BEAST 1.10 [19], which is often used for continuous phylogeographic inference.

## Results and Discussion

We here assess three distinct dispersal metrics: the weighted lineage dispersal velocity (WLDV) [20], the weighted diffusion coefficient (WDC) [21], and an IBD signal metric that we here propose to estimate as the Pearson correlation coefficient between the patristic and log-transformed great-circle geographic distances computed for each pair of tip nodes (which is similar to the metric estimated by Pekar and colleagues [22]). The great-circle distance is the geographic distance between points on the surface of the Earth and the patristic distance is the sum of the branch lengths that link two tip nodes in a phylogenetic tree. While the WLDV aims at measuring the distance covered per unit time across phylogenetic history, the WDC measures the “diffusivity” [21], i.e. the velocity at which inferred lineages invade a two-dimensional space:

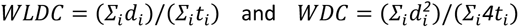

where *d*_*i*_ and *t*_*i*_ are, respectively, the great-circle distance and the time elapsed on each phylogeny branch. The concept of isolation-by-distance was first introduced by Sewall Wright [23] to describe the accumulation of local genetic differences under geographically restricted dispersal [24]. IBD analyses generally consist in determining the strength and significance of the relationship between a genetic and geographic distance between individuals [25]. The IBD signal metric considered here aims to measure to what extent phylogenetic branches are spatially structured or the tendency of phylogenetically closely related tip nodes to be sampled from geographically close locations. Note that this third metric is therefore not based on a continuous phylogeographic reconstruction but directly on a time-scaled phylogeny and sampling locations.

To evaluate the robustness of these three dispersal metrics to the sampling effort/intensity (i.e. sampling size), we have conducted phylogeographic simulations either based on a Brownian random walk (BRW) or a RRW diffusion process. For this purpose, we have implemented two distinct forward-in-time simulators, both jointly simulating time-scaled phylogenies and the dispersal history of their branches on an underlying georeferenced grid. At each time step of the BRW simulation, both the longitudinal and latitudinal displacements of evolving lineages are randomly drawn from a normal distribution; while in the RRW simulations such longitudinal and latitudinal displacements are randomly drawn from a Cauchy distribution (see the Material and Methods section for further detail). The example of a geo-referenced phylogeny simulated through a Brownian diffusion process is displayed in Figure 1 (see Figure S1 for the example of a RRW simulation). To investigate the potential impact of the phylogenetic tree inference on the different dispersal statistic estimates, we have also conducted alternative phylogeographic simulations based on tree topologies simulated under a coalescent model considering a population with a constant size (Fig. S2). The aim of all these simulations is to obtain continuous phylogeographic reconstructions that can subsequently be subsampled to artificially generate datasets that would have been obtained through various levels of sampling effort/intensity.

**Figure 1:**
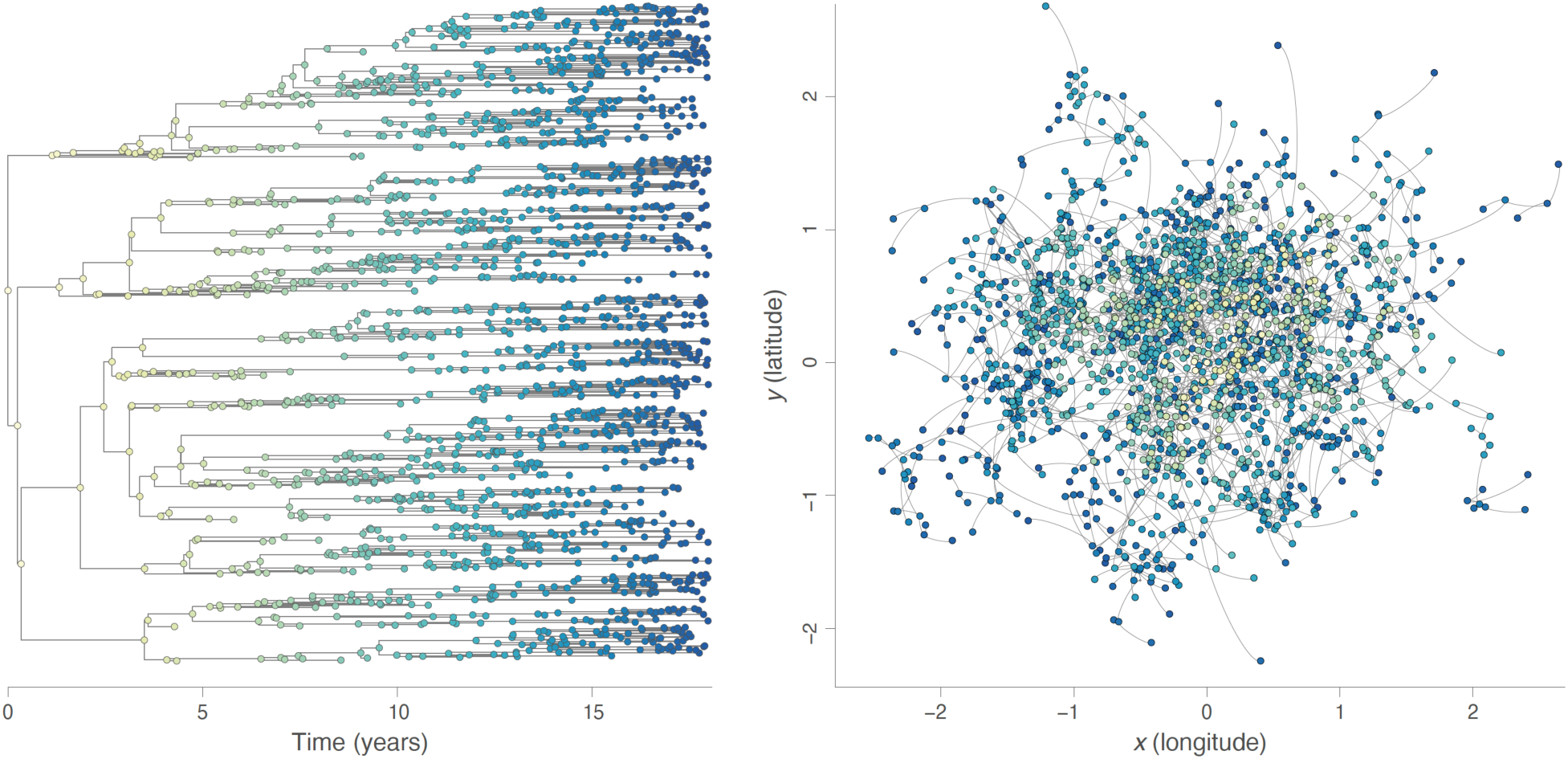
example of a continuous phylogeographic simulation based on a Brownian random walk (BRW) diffusion process. Both graphs display the phylogenetic tree sampled during a unique simulation, with its time-scaled visualisation in the left panel and its mapped visualisation in the right panel. Tree nodes are coloured according to time, with internal and tip nodes coloured according to their time of occurrence and collection time, respectively.

For each diffusion model considered, we have simulated 50 continuous phylogeographic reconstructions of more than 500 tips and then subsampled the resulting trees to only keep 500, 450, 400, 350, 300, 250, 200, 150, 100, and 50 tips (see the Material and Methods section for further detail). We have then estimated the three dispersal metrics on all the resulting datasets to explore the impact of the sampling size on their estimates (Fig. 2). Although associated with slightly more variability when estimated on the smaller datasets, both the diffusion coefficient and IBD signal metrics appear to be robust to the sampling effort (Fig. 2). This is however not the case for the lineage dispersal velocity metric for which estimates drastically decrease with the number of tips (Fig. 2). While Figure 2 reports the results obtained with the continuous phylogeographic simulations based on a Brownian diffusion process, we reach the same conclusions based on the results obtained by the simulations following a RRW diffusion process (Fig. S3) or when tree topologies are simulated under a coalescent instead of a birth-death model (Fig. S4).

**Figure 2:**
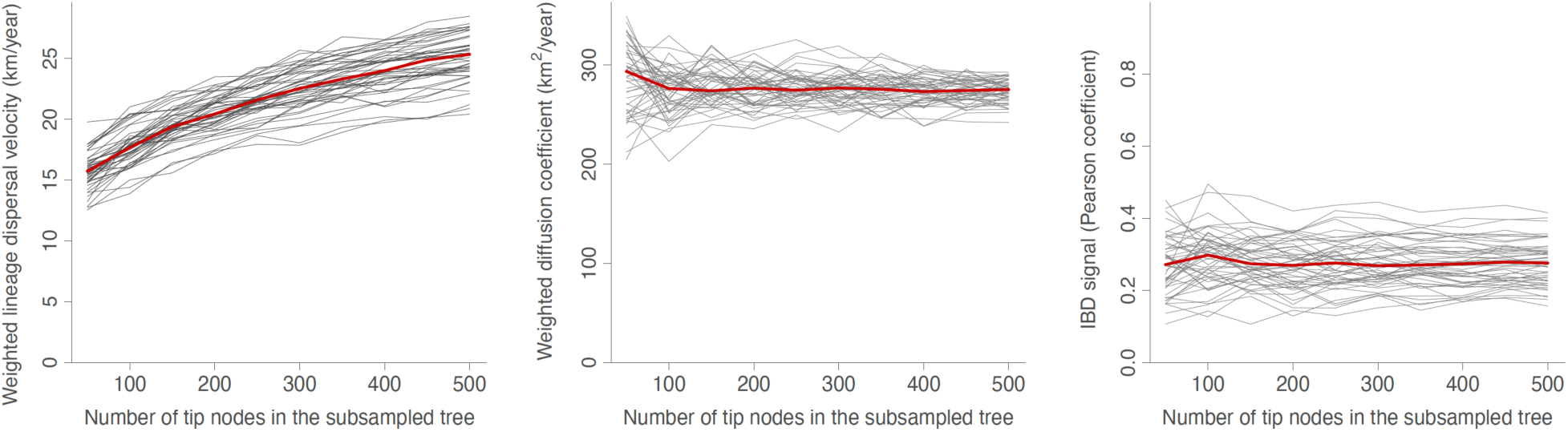
robustness of lineage dispersal metrics to the sampling effort. We here report three dispersal statistics estimated on 50 geo-referenced phylogenetic trees simulated under a Brownian diffusion process: the weighted lineage dispersal velocity (WLDV, km/year), the weighted diffusion coefficient (WDC, km^2^/year), and the isolation-by-distance (IBD) signal has been estimated by the Pearson correlation coefficient (r_P_) between the patristic and log-transformed great-circle geographic distances computed for each pair of tip nodes. Each specific tree is represented by a specific grey curve obtained when re-estimating the dispersal metric on subsampled versions of the tree, i.e. subsampled trees obtained when only randomly keeping 500, 450, 400, 350, 300, 250, 200, 150, 100, and 50 tip nodes; and the red curve indicate the median value across all simulated trees.

In addition to the three main metrics investigated above, we have also used our simulations to assess the robustness of alternative metrics, such as different ways to define the correlation coefficient measuring the IBD signal (Fig. S5) or unweighted versions of the lineage dispersal velocity and diffusion coefficient (Fig. S6). Overall, we reach the same conclusion: while the lineage dispersal velocity estimates are impacted by the sampling intensity, this does not seem to be the case for the diffusion coefficient estimates and all investigated IBD signal metrics.

From a mechanistic perspective, the lack of robustness of the lineage dispersal velocity metric can actually be expected in the context of a Brownian or, by extension, RRW diffusion process. Indeed, the sum of displacements deriving from the observation of a Brownian particle at various points in time grows with the square root of the number of (equally spaced in time) observations, making the estimation of an average speed sampling inconsistent. To illustrate this theoretical expectation, we can consider a strict one-dimensional Brownian diffusion process of position *X* along a time interval equal to 1, with *N* being the total number of regular time intervals *dt* = 1/*N, d*_*i*_ the distance travelled during time interval *i, X*_*i*_ the position at the end of interval *i*, and *X*_*pa(i)*_ the position at the end of the previous time interval. Under a strict Brownian process, we have that *X*_*i*_ – _*pa*(*i*)_ is a Gaussian random variable, so that the *L*_1_ distance covered on one interval *d*_*i*_ = |*X*_*i*_ – *X*_*pa*(*i*)_ | is distributed as a half normal variable:

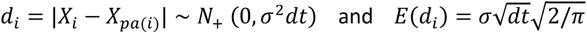

which allows to determine the expected value for the weighted lineage dispersal velocity (WLDV) as follows:

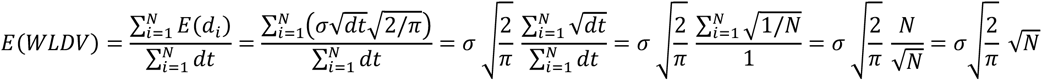

This indicates that lineage dispersal velocity depends, on average, on the number of time intervals *N* and thus the duration of these time intervals: the shorter these intervals, the larger the estimate. Because the phylogenetic branch lengths (durations) will on average decrease with the number of tip nodes, this implies, by extension, that the lineage dispersal velocity depends on the number of tip nodes included in the tree, as also illustrated by our simulations (Fig. 2). We can also follow the same reasoning to demonstrate that the expected diffusion coefficient value is not a function of *N*. Indeed, 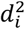 is a scaled χ^2^ random variable with one degree of freedom, so that:

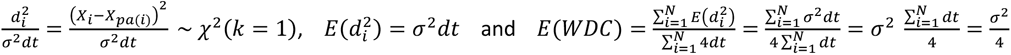

Therefore, WDC is proportional to the squared net lineage displacement per unit time (*σ*^2^) and can be used to compare the dispersal capacities of different pathogens.

In the second part of our study, we use the two dispersal metrics identified above as robust to the sampling effort — the weighted diffusion coefficient and IBD signal — to quantify and compare the dispersal capacity of viruses spreading in animal populations. We here target viruses of public and one health importance and for which comprehensive genomic datasets have at some point been exploited to conduct continuous phylogeographic reconstructions that we have managed to retrieve. The selected datasets encompass viruses associated with a broad range of host species (Table S1): moles (Nova virus), rodents (Puumala, Lassa, and Tula viruses), deer (Powassan virus), poultry and waterfowl (avian influenza virus), domestic pigs (Getah virus and porcine deltacoronavirus), cattle (lumpy skin disease virus), (migratory) bird species (West Nile virus), as well as raccoons, skunks, bats, and dogs (rabies virus). By definition, the transmission cycles of the three considered arboviruses involve transmission events through biting by infected arthropods such as ticks of the *Ixodes* and *Dermacentor* genera for the Powassan virus, as well as mosquito species of the *Culex* genus for the West Nile virus and also *Aedes* genus in the case of the Getah virus.

Our results highlight substantial differences among the datasets, both in terms of diffusion coefficient and IBD signal (Fig. 3). By far, the two virus datasets associated with the lowest diffusion coefficient are the Nova virus and Puumala virus genomic sequences collected in Belgian mole (*Talpa europaea*) and bank vole (*Myodes glareolus*) populations, respectively [26,27]. The very limited diffusion coefficient estimated for these two hantaviruses is coherent with the limited dispersal capacity of their host species. The Lassa virus dataset [28] is next in the ranking with a weighted diffusion coefficient estimated at ∼40 km^2^/year (Fig. 3, Table S1). Lassa virus is an arenavirus that also circulates in a rodent species: the Natal multimammate mouse (*Mastomys natalensis*). Interestingly, these three first datasets all reveal a relatively high IBD signal reflecting an important phylogeographic structure (Fig. 3, Table S1).

**Figure 3:**
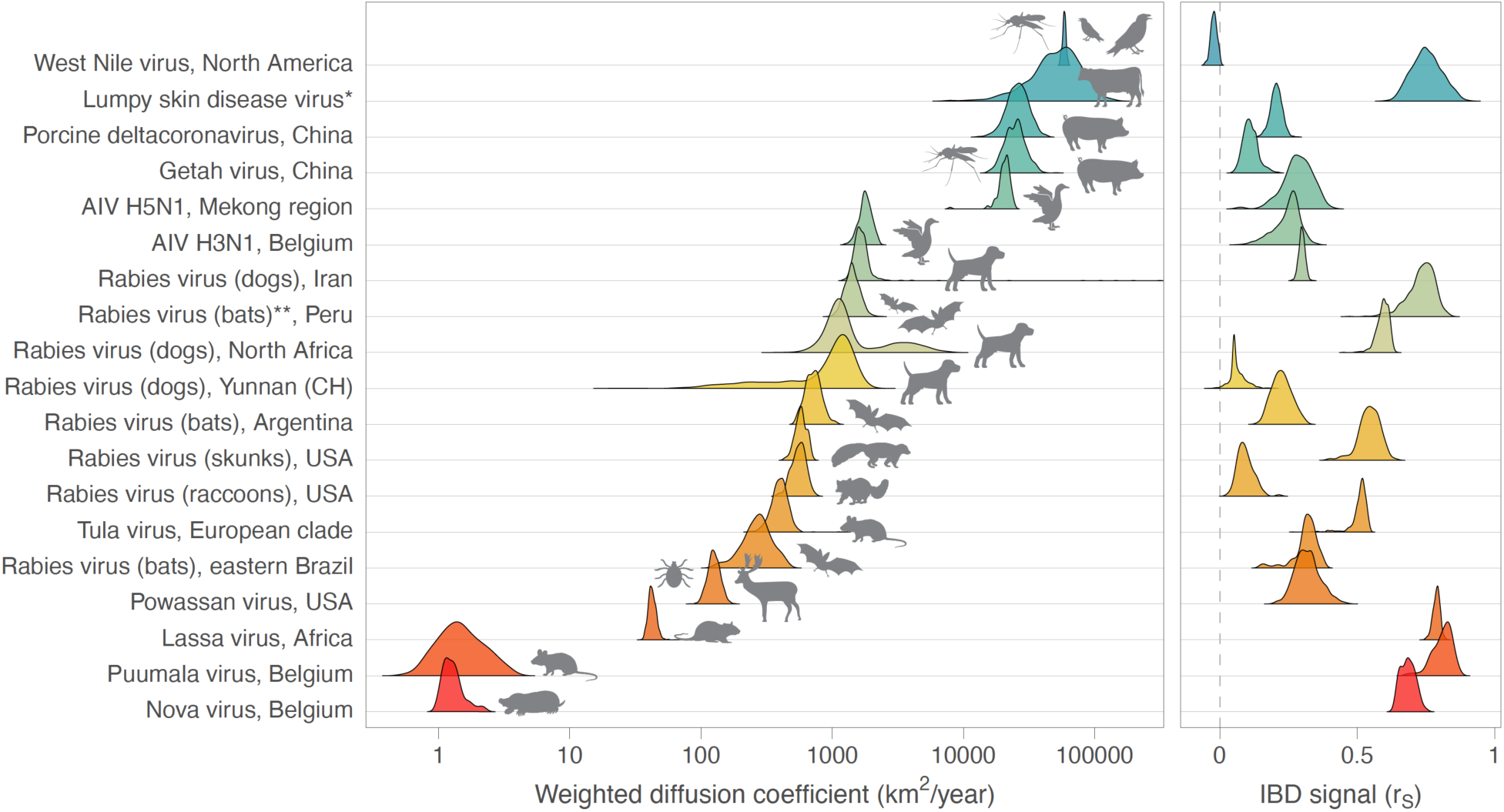
comparison of dispersal metrics estimated for different genomic datasets of viruses spreading in animal populations. Specifically, we here report posterior estimates obtained for two metrics estimated from trees sampled from the posterior distribution of a Bayesian continuous phylogeographic inference: the weighted diffusion coefficient and the isolation-by-distance (IBD) signal estimated by the Pearson correlation coefficient (r_P_) between the patristic and log-transformed great-circle geographic distances computed for each pair of virus samples. For both metrics, we report the posterior distribution of both metrics estimated through continuous phylogeographic inference for the following datasets: West Nile virus in North America [39], Lumpy skin disease virus [11], Porcine deltacoronavirus in China [40], Getah virus in China [41], avian influenza virus (AIV) in the Mekong region [42] and H3N1 in Belgium [30], rabies virus (dogs) in Iran [37], rabies virus (bats) in Peru [36], rabies virus (dogs) in northern Africa [35,44], rabies virus (bats) in Argentina [34], rabies virus (skunks) in the USA [33], rabies virus (raccoons) in the USA [32], Tula virus in central Europe [43], rabies virus (bats) in eastern Brazil [31], Powassan virus in the USA [29], Lassa virus in Africa [28], Puumala virus in Belgium [27], and Nova virus in Belgium [26]. (*) Estimates based on the analysis of the wild-type strains (see [11] for further detail); (**) estimates based on the combined analysis of lineages L1 and L3. See also Table S1 for the related 95% highest posterior density (HPD) intervals and number of samples associated with each dataset.

A large fraction of genomic datasets result in diffusion coefficient estimates between 100 and 2,000 km^2^/year (Table S1). They range from the Powassan virus dataset [29] from the United States (WDC = ∼130 km^2^/year) to the avian influenza virus (AIV) H3N1 dataset [30] corresponding to specific Belgian outbreak (∼1,800 km^2^/year). In between these two diffusion coefficient estimates, we find the diffusion coefficients estimated from a series rabies virus datasets (Fig. 3, Table S1): ∼250 km^2^/year for the bat rabies virus dataset from eastern Brazil [31], almost ∼600 km^2^/year for the raccoon [32] and skunk [33] rabies virus datasets from the USA, ∼700 km^2^/year for the bat rabies virus dataset in Argentina [34], ∼1,100 km^2^/year for the dog rabies datasets respectively from the Yunnan province (China) and northern Africa [35], ∼1,400 km^2^/year for the bat rabies dataset from Peru [36], and ∼1,600 km^2^/year for the dog/wild carnivore rabies dataset from Iran [37]. Interestingly, although the diffusion coefficient estimated for the bat rabies dataset from Peru is comparatively high, rabies viruses circulating in bats do not seem to be associated with a notably higher diffusion coefficient compared to other non-flying host species. Furthermore, it has previously been hypothesised that the relatively higher diffusion coefficient estimated for the dog rabies virus datasets compared to other rabies virus datasets could be the result of occasional rapid (long-distance) dispersal events driven by human-mediated movements [35,38]. Of note, we observe an important heterogeneity among the IBD signals estimated for the different rabies virus datasets, the IBD signal metric centred on zero for the dog rabies dataset from Yunnan or close to zero for the raccoon one from the USA, and higher than ∼0.65 for the dog rabies dataset from northern Africa and the bat rabies dataset from Peru.

We subsequently identify three data sets with diffusion coefficient estimates ranging from ∼20,000 to ∼26,000 km^2^/year including an AIV H5N1 data set from the Mekong region [42] and two genomic datasets of viruses infecting domestic pig populations in China [40,41]: the Getah virus, a re-emerging arbovirus China, and the porcine deltacoronavirus. Similarly to the hypothesis formulated for the dog rabies datasets such as the one from northern Africa, the relatively high diffusion coefficient for those two pig virus datasets may potentially be explained by long-distance dispersal through animal trade. It is also interesting to note the very limited or absent IBD signal estimated for the deltacoronavirus and Getah virus datasets, respectively, which is consistent with the notable spatial intermixing of lineage dispersal events that would be induced by animal trade across the Chinese territory [40,41].

Finally, with posterior median estimates >50,000 km^2^/year, the North American West Nile virus [39] and lumpy skin disease virus [11] datasets yield the highest diffusion coefficient estimates. Upon first detection in 1999 in New York City, the West Nile virus spread through the North American continent, carried by numerous (migratory) birds and reaching the West Coast of the United States within a period of only three years [45]. Lumpy skin disease virus was first detected in 1929 in northern Rhodesia (now Zambia) and initially circulated within the African continent. During the last decades, it has however spread eastward and to southeastern Europe [46]. Transmitted by a variety of arthropod vectors (ticks, mosquitoes, biting flies), the main hosts of this virus are cattle and buffalos while the role of other wild animals in its transmission and spread remains uncertain [46]. The continuous phylogeographic reconstruction of the lumpy skin disease virus dissemination considered here is a global one involving long-distance lineage dispersal events among continents. Also in this case, the high diffusion coefficient estimated for this dataset is potentially related to the transport of infected cattle to naive regions [11]. It is also worth noting that the IBD signal estimated for the West Nile virus and lumpy skin disease virus datasets are drastically different, being centred on zero for the former and approximately 0.8 for the latter. This result really reflects the phylogeographic patterns reconstructed for both outbreaks, the West Nile virus one displaying an important amount of longitudinal, and to some extent latitudinal, back and forth lineage dispersal events during the endemic phase of the North American outbreak, and the lumpy skin disease virus spread mainly consisting in long-distance dispersal events establishing more local transmission chains.

Furthermore, the relatively important diffusion coefficient estimated for the WNV dataset appears to be partially due to the expansion phase of this outbreak: when computing a distinct diffusion coefficient for the expansion and endemic phases — here approximated by considering phylogenetic branches occurring before or after 2004 [39], respectively — we obtain an estimate that is between two and three times higher for the expansion phase (Table S1), illustrating the impact of the distinction of those epidemic phases on the metric. In the case of the WNV spread across the North American continent, the invasion was in part driven by long-distance dispersal events, which can transform the spread of a pathogen from a wave-like to a fast “metastatic” growth pattern [47]. Overall, the comparison of the estimates based on the WNV dataset highlights that a metric like the diffusion coefficient can notably be impacted by the epidemic phase of the considered outbreak, an expansion phase being more likely to be associated with higher estimates than an endemic phase.

### Limitations, Conclusions, and Perspectives

Our study comes with a number of limitations. First, while our simulation framework explores the impact of sampling intensity (i.e. heterogeneity in the *absolute* sampling effort) on dispersal statistic estimates, it does not, however, include varying degrees of sampling bias (i.e. *spatial* heterogeneity in the sampling effort) nor border effects, which can also affect the target statistics. For instance, Layan and colleagues have illustrated that the extent of the study area can impact the diffusion coefficient estimates [48]. As an interpretation, they put forward that “larger sampling areas are expected to be associated with higher probabilities to sample long-distance dispersal events that will, on average, more likely correspond to fast dispersal events than short-distance dispersal events” [48]. As detailed in Table S1, we however see that the different datasets analysed here encompass a large range of study area sizes that are not necessarily following the trend of diffusion coefficient estimates. Overall, the impact of the spatial heterogeneity in the sampling effort on the outcomes of phylogeographic reconstructions has previously been assessed and discussed by Kalkauskas and colleagues [49].

Second, our simulation framework does not involve the generation of actual genomic sequences. Because the lineage dispersal metrics investigated here are directly estimated from the phylogenetic trees simulated on a map, we do not evaluate to what extent the phylogenetic errors will contribute to the estimation uncertainty of these statistics. From a computational perspective, adding a genomic sequence simulation step would have also drastically increased the computation time and resources required to conduct the analyses. This would indeed imply running no less than one thousand continuous phylogeographic analyses (corresponding to both Brownian and RRW simulations, times fifty simulations, times the generation of ten subsampled datasets with a decreasing number of tip nodes).

Spatially explicit phylogeographic reconstructions are relatively widely applied thanks to popular implementations of the RRW model [7,15,50], but there is a need for robust metrics allowing a direct comparison of lineage dispersal dynamics for different viral variants/outbreaks or even distinct viruses. Our simulations highlight that, among the three different kinds of dispersal metrics investigated here, only two of them appear to be robust to the sampling intensity: the diffusion coefficient and the IBD signal metrics, the first one being estimated from continuous phylogeographic inference and the second one being directly estimated from a (time-scaled) phylogeny and sampling locations. The lineage dispersal velocity estimated from continuous phylogeographic inference on the other hand increases with the number of samples in the phylogeographic analysis. We subsequently confirmed that this trend can be explained by a mechanistic dependence of this metric on the number of genomic sequences (tip nodes) considered in the analysis.

The two metrics identified as robust are measuring complementary aspects of the overall dispersal pattern: while the diffusion coefficient allows estimating a diffusion coefficient as the invaded area per unit of time, the IBD signal metric will quantify to what extended the geo-referenced phylogenies are spatially structured, i.e. the tendency for phylogenetically-related lineages to evolve in a close vicinity. In terms of readability, the diffusion coefficient (WDC) can also be used to estimate the quadratic mean — also called root mean square — of the effective displacement by unit of time, which is equal to 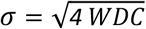. Furthermore, if the mean time needed for a pathogen to infect successive hosts by contagion (a kind of generation time, *T*) was known, the root mean square dispersal distance between two successive hosts could be estimated as 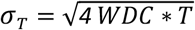.

In the second part of our study, we have taken advantage of these two robust metrics to compare the dispersal pattern of viral lineages informed by the available continuous phylogeographic reconstruction of a large range of published datasets of genomic sequences. Our results confirm and highlight an important heterogeneity in the dispersal capacity of viruses primarily spreading in animal populations, directly reflecting the dispersal capacity and/or the human-mediated movement of the main host species carrying those viruses. The comparative framework initiated in the present study will allow extending the comparison of virus dispersal capacities to new genomic datasets analysed through continuous phylogeographic inference. In practice, the GitHub repository referenced below can be used to add additional datasets in the comparison reported in Figure 3 and Table S1, with the perspective to further unveil and understand the velocity at which viruses can circulate in various host populations.

In our view, the field of phylogeography, or phylodynamic in general, applied to fast-evolving pathogens such as RNA (and DNA) viruses faces a series of (new) challenges. A first challenge lies in the recent and impressive increase in the amount of genomic information that can be exploited in phylodynamic analyses. Driven by technological developments in next-generation sequencing [54–56] and important national budgets unlocked for genomic surveillance in the context of the COVID-19 pandemic, the number of geo-referenced genomic sequences available for conducting phylogeographic reconstruction of some viruses of public health importance has exploded, which is particularly true for SARS-CoV-2 with >16.5 million full genomes available on the GISAID platform to date (March 2024, https://gisaid.org/). Unfortunately, fully Bayesian phylogeographic approaches reach their computational limit when analysing datasets consisting of a few thousand genomes. While recent developments have been dedicated to the phylogenetic inference of large-scale genomic datasets (see e.g. [57,58]), facing computational limitations will continue to be a timely challenge in the development of phylodynamic approaches aiming to look below the tip of the iceberg by exploiting the large amount of pathogen genetic information at our disposal in order to get quantitative insights into or test hypotheses on ongoing epidemics. Another challenge likely lies in the perspective to go beyond the retrospective aspect of phylodynamic analyses and feed the knowledge obtained through such reconstructions into predictive frameworks.

## Materials and Methods

### Continuous phylogeographic simulations

We conducted continuous phylogeographic simulations based on both a Brownian (BRW) and a relaxed (RRW) random walk diffusion process on an underlying georeferenced grid (raster) with a resolution of 0.5 arcmin. We first implemented a forward-in-time birth-death approach to simulate the dispersal history of viral lineages over twenty years while considering a time step of one day. Specifically, each simulation started from an index case placed in the centre of the grid. At each time step, each ongoing lineage had (i) a first probability to give birth to another lineage (*p*_birth_ = 0.6 per year, so 0.0016 per day) and (ii) a second probability to be “sampled” (*p*_sampling_ = 0.4 per year, so 0.0011 per day), i.e. to stop evolving and leave the simulation. Of note, we defined a sampling window so that evolving lineages could not be sampled during the first two years of the simulation/outbreak. In between the two, ongoing lineages also had the opportunity, at each time step, to move on the grid. In the BRW simulations, the longitudinal and latitudinal displacements were stochastically defined by respectively drawing an horizontal *d*_x_ and vertical *d*_y_ displacement value in a normal distribution of mean equal to zero and standard deviation here arbitrarily set to 0.0125; and in the RRW simulations, the longitudinal and latitudinal displacements were stochastically defined by respectively drawing an horizontal *d*_x_ and vertical *d*_y_ displacement value in a Cauchy distribution with a position parameter set to zero and a scale parameter set to 2.5*x*10^-4^. In both cases, the resulting displacement on the map was then randomly rotated around the previous location point. In particular for the RRW simulations, this stochastic rotation step prevents rare long-distance displacement to only occur along a latitudinal or longitudinal gradient. Overall, we conducted 200 BRW and 200 RRW simulations and retained in each case the first 50 simulations leading to simulated phylogenies with at least 500 tip nodes. Selected phylogeographic simulations were then eventually subsampled to obtain georeferenced time-scaled phylogenies only including 500, 450, 400, 350, 300, 250, 200, 150, 100, and 50 tip nodes. The script used to conduct the BRW and RRW simulations was implemented in a new function named “simulatorRRW2” and included in the R package “seraphim” [20,59] (the “simulatorRRW1” function corresponding to another simulator previously implemented to conduct simulations along tree topologies [60]).

In addition to the simulations with phylogenetic trees simulated under a birth-death model, we also conduct BRW simulations along trees primarily simulated under a coalescent model. To perform these coalescent simulations, we used the function “rcoal” of the R package “ape”. Longitudinal and latitudinal displacements were then simulated along the resulting tree topologies with the “fastBM” function of the R package “phytools” [61]. As for the simulations based on a birth-death model, we conducted 200 simulations and retained in each case the first 50 simulations leading to simulated phylogenies with at least 500 tip nodes, and those phylogeographic simulations were eventually subsampled to obtain georeferenced time-scaled phylogenies only including 500, 450, 400, 350, 300, 250, 200, 150, 100, and 50 tip nodes.

### Dispersal statistics estimation

Dispersal statistics were all computed using the “spreadStatistics” function in the R package “seraphim” [20,59], which now also allows the estimation of the IBD signal computed as the Pearson correlation (r_P_) between the patristic and log-transformed great-circle geographic distances computed for each pair of virus samples. 95% highest posterior density (HPD) intervals were computed with the “hdi” function of the R package “HDInterval”, and the ridgeline plots were generated using the “ridgeline” R package. As mentioned above, in complement to the weighted lineage dispersal velocity (WLDV) and weighted diffusion coefficient (WDC) metrics, we also computed and tested unweighted versions — the mean lineage dispersal velocity (MLDV) [20] and mean diffusion coefficient (MDC) [15]:

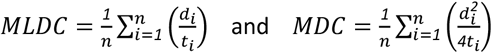

where *n* is the number of phylogenetic branches, *d*_*i*_ the geographic (great-circle) distance, and *t*_*i*_ the time elapsed on each phylogeny branch.

## Code and data availability

R scripts related to the analyses based on simulated and real datasets are all available, along with the associated input/output files, at https://github.com/sdellicour/dispersal_capacities. Continuous phylogeographic simulations and dispersal statistics were respectively conducted and computed using the R package “seraphim” available at https://github.com/sdellicour/seraphim (see also the updated “seraphim” tutorial on the estimation of dispersal statistics available at : https://github.com/sdellicour/seraphim/blob/master/tutorials/).

## Acknowledgements

We are grateful to Richard Neher for his useful feedback on this study. SD and PL acknowledge support from the European Union Horizon 2020 project MOOD (grant agreement n°874850). SD also acknowledges support from the *Fonds National de la Recherche Scientifique* (F.R.S.-FNRS, Belgium; grant n°F.4515.22), from the Research Foundation — Flanders (*Fonds voor Wetenschappelijk Onderzoek — Vlaanderen*, FWO, Belgium; grant n°G098321N), and from the European Union Horizon 2020 project LEAPS (grant agreement n°101094685). PR’s internship at the University of Montpellier was founded by the I-SITE MUSE through the Key Initiative “Data and Life Sciences”. MAS and PL acknowledge support from the European Union’s Horizon 2020 research and innovation programme (grant agreement no. 725422-ReservoirDOCS), from the Wellcome Trust through project 206298/Z/17/Z, and from the National Institutes of Health grants R01 AI153044, R01 AI162611 and U19 AI135995. PL also acknowledges support from the Research Foundation — Flanders (*Fonds voor Wetenschappelijk Onderzoek — Vlaanderen*, FWO, Belgium; grants n°G0D5117N and G051322N).

---

**Figure S1:**
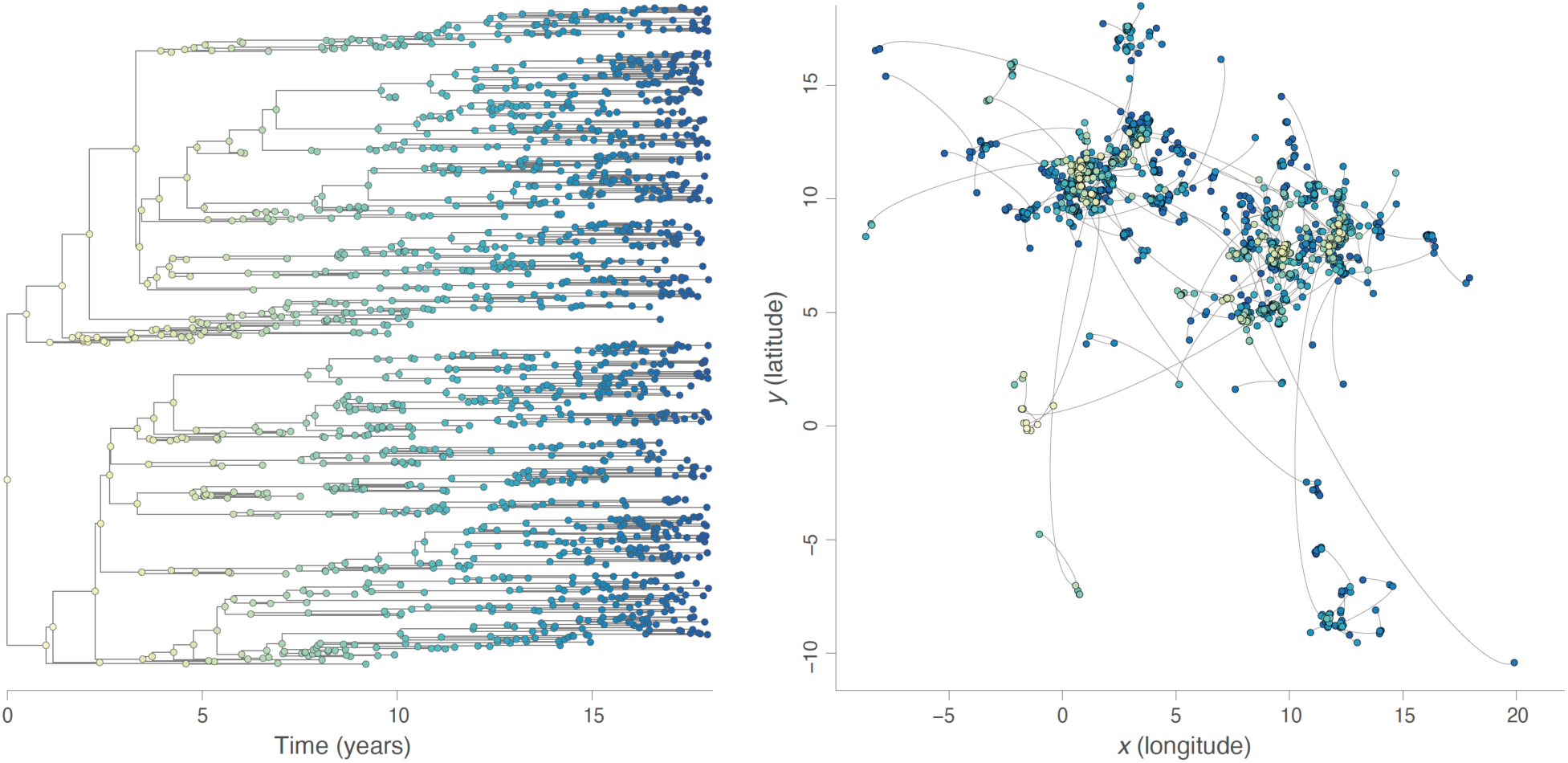
example of a continuous phylogeographic simulation based on a relaxed random walk (RRW) diffusion process with phylogenetic trees simulated under a birth-death model. Both graphs display the phylogenetic tree sampled during a unique simulation, with its time-scaled visualisation in the left panel and its mapped visualisation in the right panel. Tree nodes are coloured according to time, with internal and tip nodes coloured according to their time of occurrence and collection time, respectively.

**Figure S2:**
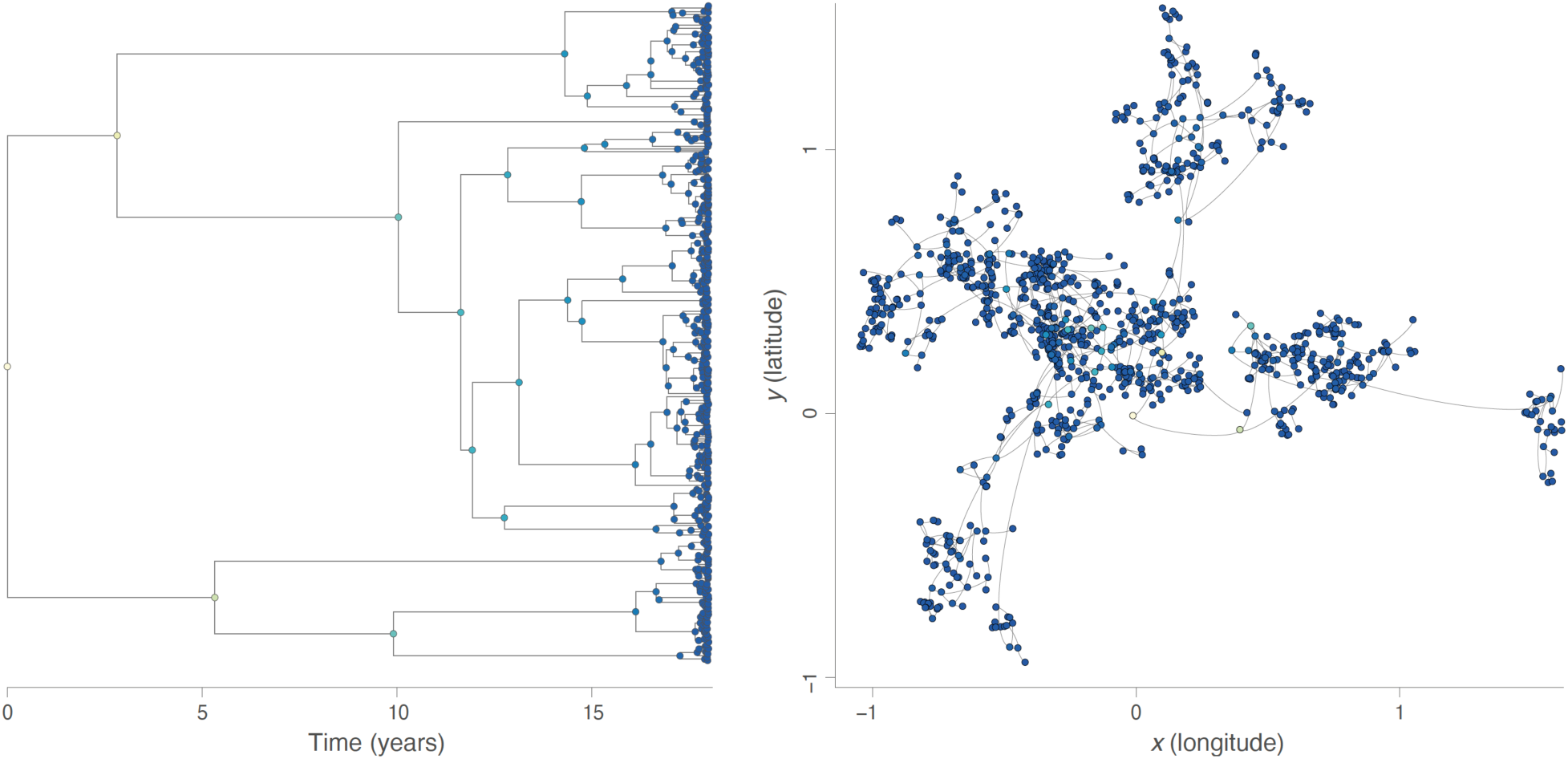
example of a continuous phylogeographic simulation based on a relaxed random walk (RRW) diffusion process with phylogenetic trees simulated under a coalescent model. Both graphs display the phylogenetic tree sampled during a unique simulation, with its time-scaled visualisation in the left panel and its mapped visualisation in the right panel. Tree nodes are coloured according to time, with internal and tip nodes coloured according to their time of occurrence and collection time, respectively.

**Figure S3:**
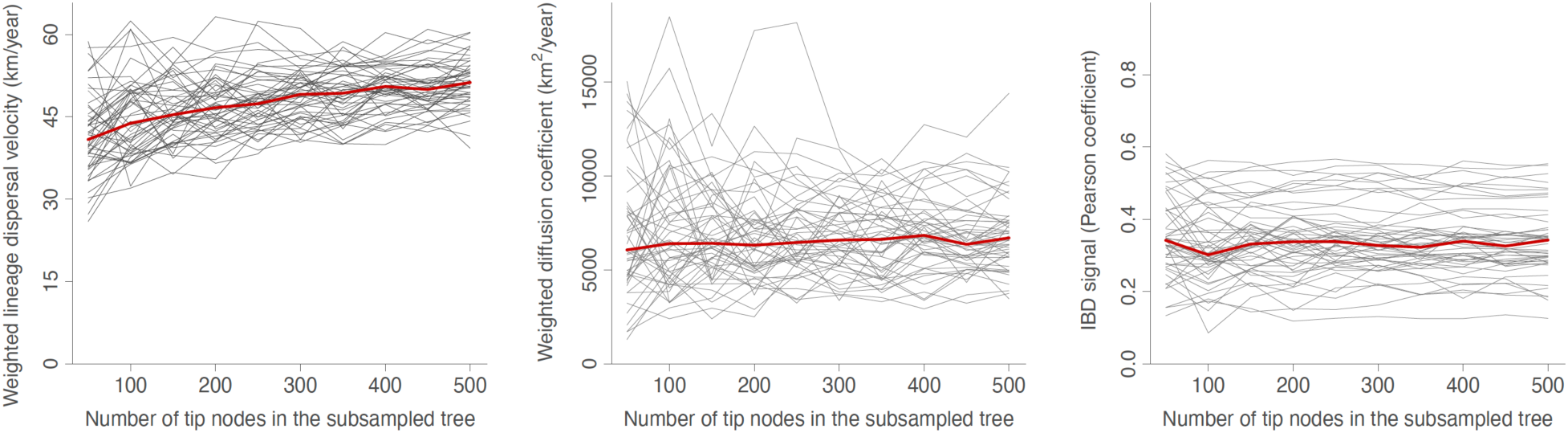
robustness of lineage dispersal metrics to the sampling effort, here based on the relaxed random walk (RRW) simulations where phylogenetic trees were simulated under a birth-death model. In complement to Figure 1 reporting the corresponding results based on Brownian random walk (BRW) simulations, we here report the same dispersal statistics estimated on 50 geo-referenced phylogenetic trees simulated under a RRW diffusion process: the weighted lineage dispersal velocity (km/year), the weighted diffusion coefficient (km^2^/year), and the isolation-by-distance (IBD) signal estimated by the Pearson correlation coefficient between the patristic and log-transformed great-circle geographic distances computed for each pair of tip nodes. Each specific tree is represented by a specific grey curve obtained when re-estimating the dispersal metric on subsampled versions of the tree, i.e. subsampled trees obtained when only randomly keeping 500, 450, 400, 350, 300, 250, 200, 150, 100, and 50 tip nodes; and the red curve indicate the median value across all simulated trees.

**Figure S4:**
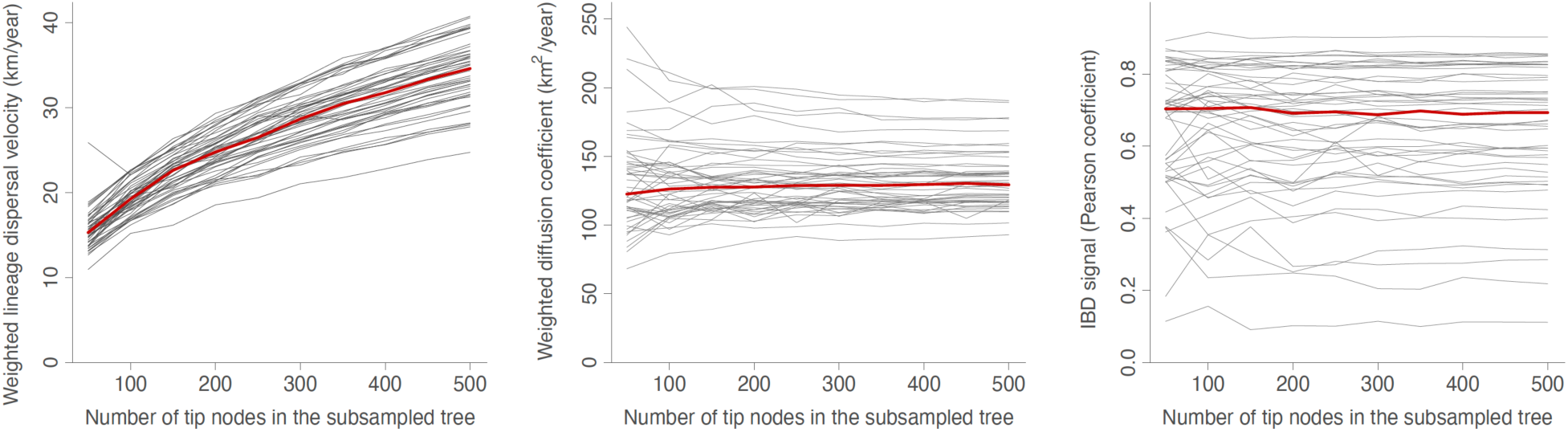
robustness of lineage dispersal metrics to the sampling effort, here based on the Brownian random walk (BRW) simulations where phylogenetic trees were simulated under a coalescent model. In complement to Figure 1 reporting the corresponding results from BRW simulations where phylogenetic trees were simulated under a birth-death model, we here report the same dispersal statistics estimated on 50 geo-referenced phylogenetic trees simulated under a BRW diffusion process this time based on tree topologies simulated under a coalescent model: the weighted lineage dispersal velocity (km/year), the weighted diffusion coefficient (km^2^/year), and the isolation-by-distance (IBD) signal has been estimated by the Pearson correlation coefficient between the patristic and log-transformed great-circle geographic distances computed for each pair of tip nodes. Each specific tree is represented by a specific grey curve obtained when re-estimating the dispersal metric on subsampled versions of the tree, i.e. subsampled trees obtained when only randomly keeping 500, 450, 400, 350, 300, 250, 200, 150, 100, and 50 tip nodes; and the red curve indicate the median value across all simulated trees.

**Figure S5:**
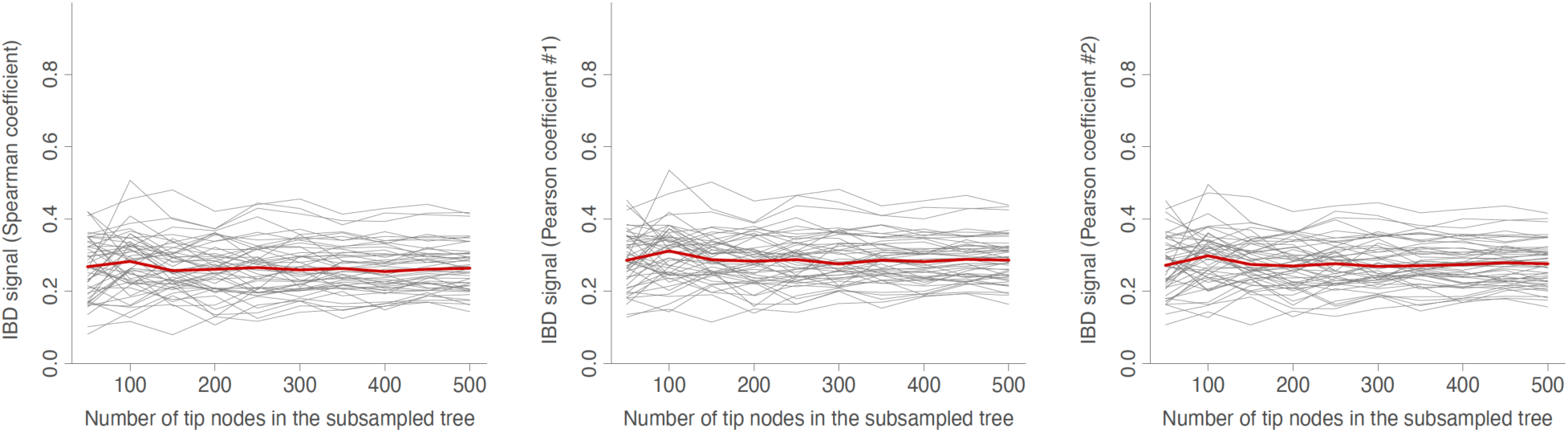
robustness of alternative isolation-by-distance (IBD) metrics to the sampling effort, here based on the Brownian random walk (BRW) simulations where phylogenetic trees were simulated under a birth-death model. In complement to Figure 1, we here report three distinct iBD signal metrics estimated on the same 50 geo-referenced phylogenetic: (i) the Spearman correlation coefficient between the patristic and great-circle geographic distances computed for each pair of tip nodes, (ii) the Pearson correlation coefficient (#1) between the patristic and great-circle geographic distances computed for each pair of tip nodes, and (ii) the Pearson correlation coefficient (#2) between the patristic and log-transformed great-circle geographic distances computed for each pair of tip nodes. Each specific tree is represented by a specific grey curve obtained when re-estimating the dispersal metric on subsampled versions of the tree, i.e. subsampled trees obtained when only randomly keeping 500, 450, 400, 350, 300, 250, 200, 150, 100, and 50 tip nodes; and the red curve indicate the median values.

**Figure S6:**
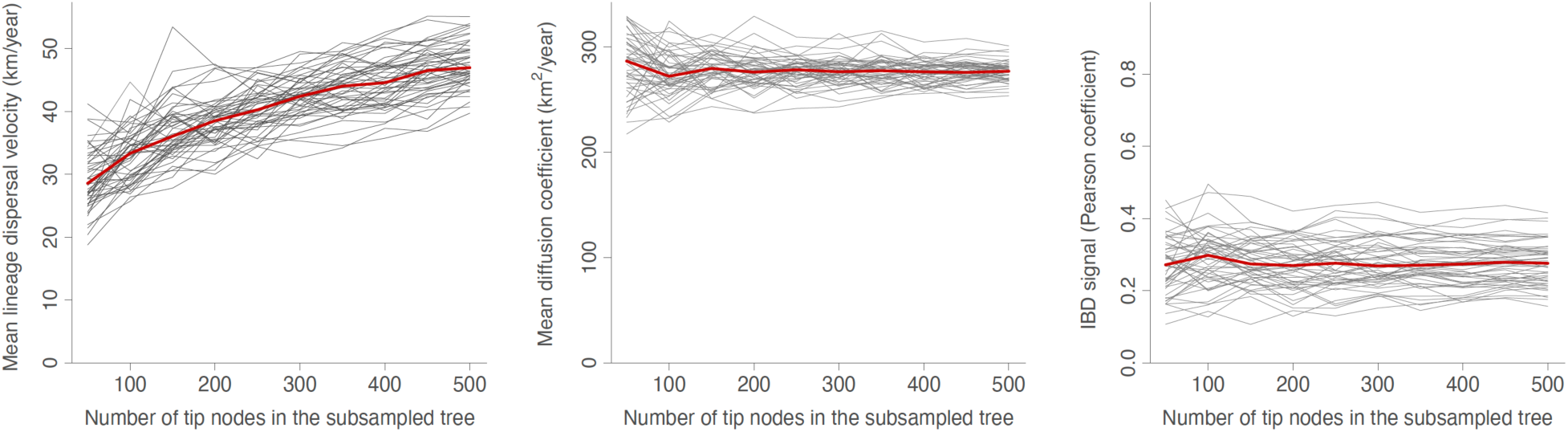
robustness of alternative dispersal metrics to the sampling effort, here based on the Brownian random walk (BRW) simulations where phylogenetic trees were simulated under a birth-death model. In complement to Figure 1 reporting the results obtained for the weighted lineage dispersal velocity (WLDV, km/year) and weighted diffusion coefficient (WDC, km^2^/year), we here report the mean lineage dispersal velocity (MLDV, km/year) and mean diffusion coefficient (MDC, km^2^/year) estimates. As in Figure 1, we also report estimates for the isolation-by-distance (IBD) signal has been estimated by the Pearson correlation coefficient between the patristic and log-transformed great-circle geographic distances computed for each pair of tip nodes. Each of the 50 simulated trees is represented by a specific grey curve obtained when re-estimating the dispersal metric on subsampled versions of the tree, i.e. subsampled trees obtained when only randomly keeping 500, 450, 400, 350, 300, 250, 200, 150, 100, and 50 tip nodes; and the red curve indicate the median value across all simulated trees.

**Table S1:** comparison of dispersal metrics estimated for different genomic datasets of viruses spreading in animal populations. The isolation-by-distance (IBD) signal has been estimated by the Pearson correlation coefficient (r_P_) between the patristic and log-transformed great-circle geographic distances computed for each pair of virus samples. For each dataset and metric, we report both the posterior median estimate and the 95% highest posterior density (HPD) interval. The extent of the study areas were approximated by the inland area of the minimum convex hull polygon surrounding all the sampling locations of a given dataset. “AIV” refers to avian influenza virus; (*) estimates based on the combined analysis of lineages L1 and L3; (**) estimates based on the analysis of the wild-type strains (see [11] for further detail); (***) different weighted diffusion coefficient estimates are obtained when solely focusing on phylogeny branches occurring before (113918 km^2^/year, 95% HPD = [102250, 136202]) and after (43400 km^2^/year, 95% HPD = [38943, 49230]) 2004; which approximately corresponds to the end of what can be characterised as the expansion phase of the WNV outbreak in North America.

| Dataset | Weighted diffusion coefficient (km<sup>2</sup>/year) | IBD signal ($r_p$) | Num. of samples | Study area size (km<sup>2</sup>) | Reference |
| --- | --- | --- | --- | --- | --- |
| Nova virus, Belgium | 1 [1, 2] | 0.68 [0.64, 0.73] | 100 | ~8,000 | Laenen *et al.* (2016) |
| Puumala virus, Belgium | 1 [1, 3] | 0.82 [0.75, 0.87] | 66 | ~8,000 | Laenen *et al.* (2019) |
| Lassa virus, segment S, Africa | 42 [38, 49] | 0.79 [0.76, 0.81] | 410 | ~1,255,000 | Klitting *et al.* (2022) |
| Powassan virus, USA | 126 [101, 154] | 0.32 [0.24, 0.42] | 273 | ~118,000 | Vogels *et al.* (2023) |
| Rabies virus (bats), eastern Brazil | 274 [149, 427] | 0.32 [0.20, 0.38] | 41 | ~44,000 | Vieira *et al.* (2013) |
| Tula virus, European clade | 399 [282, 506] | 0.52 [0.45, 0.55] | 380 | ~1,270,000 | Cirkovic *et al.* (2022) |
| Rabies virus (raccoons), USA | 561 [453, 706] | 0.09 [0.05, 0.16] | 47 | ~457,000 | Biek *et al.* (2007) |
| Rabies virus (skunks), USA | 580 [477, 679] | 0.55 [0.46, 0.61] | 241 | ~1,892,000 | Kuzmina *et al.* (2013) |
| Rabies virus (bats), Argentina | 721 [550, 909] | 0.22 [0.16, 0.29] | 131 | ~142,000 | Torres *et al.* (2014) |
| Rabies virus (dogs), Yunnan (CH) | 1064 [75, 1544] | 0.06 [0.01, 0.13] | 247 | ~301,000 | Tian *et al.* (2018) |
| Rabies virus (dogs), North Africa | 1191 [546, 4844] | 0.60 [0.55, 0.63] | 250 | ~667,000 | Talbi *et al.* (2010) |
| Rabies virus (bats)*, Peru | 1416 [1086, 1812] | 0.74 [0.64, 0.82] | 260 | ~675,000 | Streicker *et al.* (2016) |
| Rabies virus (dogs), Iran | 1643 [1307, 2135] | 0.30 [0.28, 0.32] | 105 | ~1,052,000 | Dellicour *et al.* (2019) |
| AIV H3N1, Belgium | 1794 [1414, 2200] | 0.26 [0.14, 0.33] | 101 | ~5,000 | Van Borm *et al.* (2023a) |
| AIV H5N1, Mekong region | 20647 [16951, 24285] | 0.29 [0.18, 0.37] | 320 | ~956,000 | Dellicour *et al.* (2020a) |
| Getah virus, China | 24930 [17065, 34887] | 0.11 [0.06, 0.17] | 125 | ~4,060,000 | Zhao *et al.* (2023) |
| Porcine deltacoronavirus, China | 26093 [16753, 36264] | 0.21 [0.16, 0.25] | 97 | ~1,967,000 | He *et al.* (2020) |
| Lumpy skin disease virus** | 52989 [16749, 99297] | 0.75 [0.66, 0.86] | 34 | ~31,658,000 | Van Borm *et al.* (2023b) |
| West Nile virus, North America*** | 58757 [55087, 62105] | -0.02 [-0.04, 0.00] | 801 | ~7,947,000 | Dellicour *et al.* (2020b) |

